# Old Collections, New Perspectives: Analytical Reassessment of One of Europe’s Biggest Archaeological Broomcorn Millet Deposits (Romania)

**DOI:** 10.1101/2025.11.26.690670

**Authors:** Ana García-Vázquez, Mihaela Golea, Cristina Mircea, Marin Cârciumaru, Gabriela Sava, Ana Ilie, Cătălin Lazăr

## Abstract

The broomcorn millet grains (*Panicum miliaceum*) analysed in this study, recovered from Morteni tell (Romania), represent one of Europe’s largest archaeological millet deposits (150 kg). Originally attributed to the Chalcolithic period (Gumelniţa culture), radiocarbon dating revealed a significantly later context (361 cal BC – 61 cal AD (95.4%)), placing the grains in the Late Iron Age - Latène, when millet was an established staple in the region. The uncharred, desiccated grains exhibited uneven preservation: while some retained isotopic values akin to fresh millet, others showed evidence of nitrogen loss due to diagenesis. High δ^15^N values point to intensive manuring, highlighting millet’s agricultural significance during the Iron Age. Combining isotopic data with FTIR-ATR analysis allowed the detection of diagenetic impacts and underscored the importance of cross-disciplinary approaches in archaeobotanical studies. These findings demonstrate the necessity of radiocarbon dating and integrated analyses to avoid misinterpretations based on stratigraphy alone and to enhance the reliability of data on ancient crop use and significance.

## 1. Introduction

Millets, small-seeded cereal crops of the Poaceae family, are highly valued for their resilience in arid and semi-arid regions, thriving under drought and poor soil conditions. As C4 plants, millets utilise an efficient photosynthetic pathway, enabling high carbon fixation even in high-temperature and low-moisture environments. Their short growing season of up to three months allows for an additional summer harvest, making them a critical safeguard against famine when winter crops fail. Among the most prominent millets, broomcorn millet (*Panicum miliaceum*) and foxtail millet (*Setaria italica*) were domesticated in northern China as early as 10,300–8,700 cal BP (Lu et al. 2009; Zhao 2011; Liu et al. 2012; Yang et al. 2012; Wang et al. 2017), spreading widely due to their adaptability. However, radiocarbon dating has revealed that millet cultivation in Europe began significantly later, with evidence suggesting its introduction around the 16^th^ century BC and rapid spread during the 15^th^ and 14^th^ centuries BC (Motuzaite-Matuzeviciute et al. 2013; Filipović et al. 2020).

In Romania (Supplementary Table 1), archaeobotanical evidence of broomcorn millet has been identified at Neolithic and Chalcolithic sites, though these finds are often isolated and lack radiocarbon confirmation (Cârciumaru 1996; Daisă-Ciută et al. 2004; Monah and Monah 2005; Ciută 2006; Monah 2007; Lazăr et al. 2020), raising concerns about their reliability. This is the case with broomcorn millet grains found at Măgura-Buduiasca, originally attributed to Neolithic occupation levels associated with the Dudesti culture (6^th^ millennium BC) (Bogaard and Walker 2011). Radiocarbon dating, however, revealed much later dates, ranging from 1434-1268 cal BC and 1438-1620 cal AD (Motuzaite-Matuzeviciute et al. 2013). Similarly, the broomcorn millet remains from Dobrovăţ and Baia-În Muchie, originally attributed to the Cucuteni A and Pre-Cucuteni levels, yielded significantly more recent dates (An et al. 2025). Millet cultivation became more prominent in the Bronze Age and widespread by the Iron Age, appearing frequently at Dacian sites (Cârciumaru 1983; Cârciumaru 1996). Historical references to millet in the region are scarce, with Herodotus noting its use among the Scythians, potentially linked to the Dacians. Millet was a dietary staple until the 17^th^ century AD, when corn (*Zea mays*) replaced it (Panţu 1906). Today, its use is limited, with *braga* or *boza*, a sweet, fermented drink of Turkish origin, being one of the few traditional recipes preserved (Roman 1998).

Continuing the recent efforts to revisit archaeobotanical archives from prehistoric sites in Romania (Golea et al. 2023), this study re-examines one of the largest archaeological millet deposits (150 kg) discovered at the Chalcolithic tell site of Morteni (Gumelniţa culture). Initially attributed to the Chalcolithic period (Diaconescu 1978-1979; Cârciumaru 1996), this assignment has since been recognised as inaccurate. The misclassification reflects the methodological limitations of archaeobotanical research in the 1970s–1980s, which often associated millet usage indiscriminately with prehistoric communities. To address these inaccuracies, perpetuated in subsequent studies (Ilie and Dumitru 2010; Golea 2016), this paper integrates radiocarbon dating, isotopic analyses, and 3D morphometric techniques to reassess the Morteni deposit. This comprehensive approach provides a revised chronological framework and cultural context, clarifying the significance of one of Europe’s largest archaeological millet deposits and contributing to a more accurate understanding of millet’s role in prehistoric subsistence strategies.

## 2. Material and Methods

### 2.1. The site

The Morteni site, known as “Măgura” or “La Movilă”, is located on the western edge of the village of the same name in Dâmboviţa County, Romania (Lat. EPSG_384: 517665.358 / Long. EPSG_384: 352285.353) in northen Balkans (Figure 1A). This tell-type settlement appears as a circular mound that covers an area of approximately 5,000 m^2^ and reaches a maximum elevation of 197 m.a.s.l., standing 3 to 5 m above the current ground level. The site is bordered by the Măgura River, a tributary of the Neajlov, which flows about 2 km away (Figure 1B). Unfortunately, the site is not protected, as it is situated in an agricultural area, and is constantly subjected to ploughing (Diaconescu 1978-1979; Ilie and Dumitru 2010; Ştefan 2010; Covătaru 2024).

**Figure 1.**
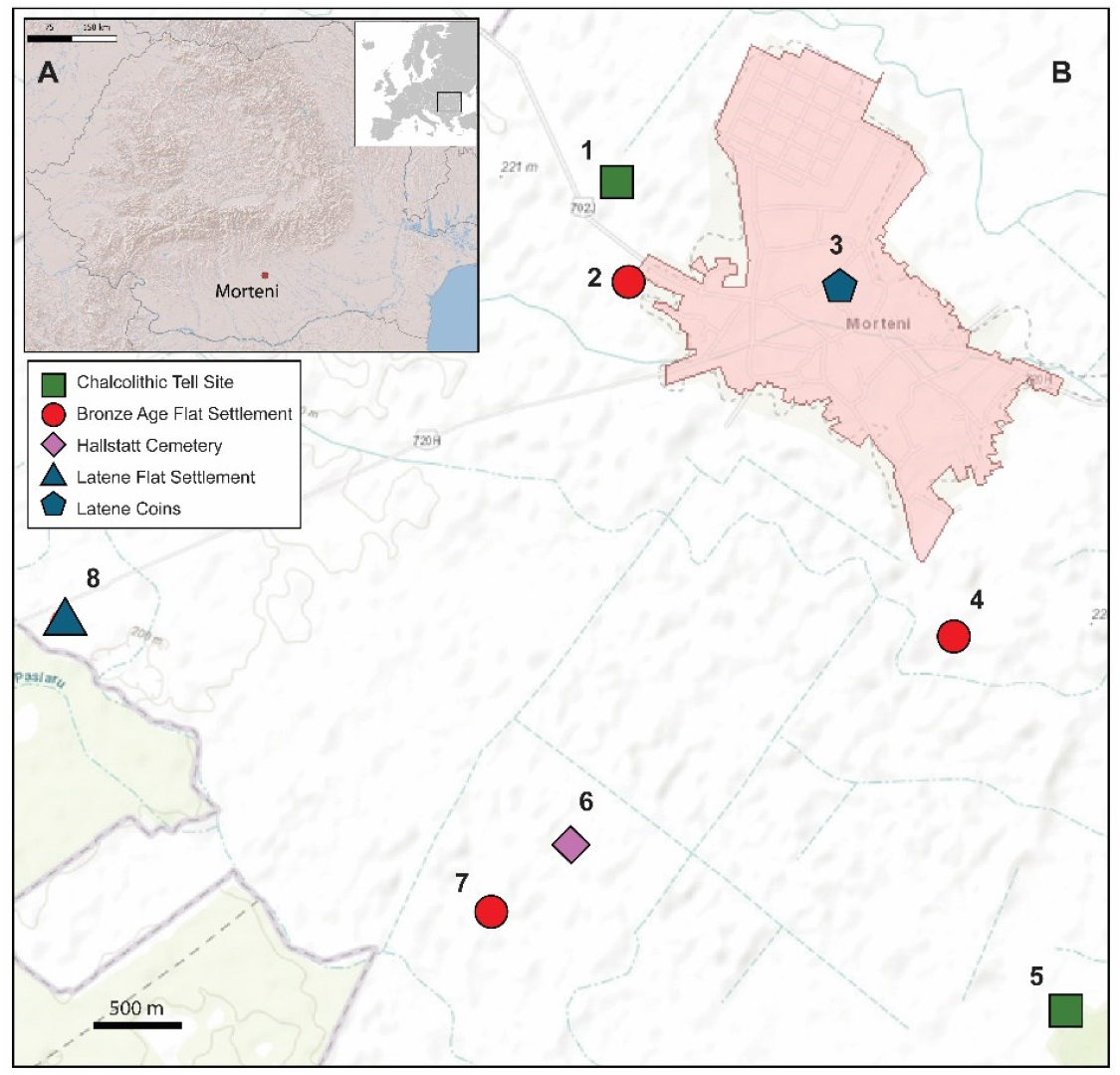
(A) Geographical location of the Morteni site. (B) Placement of the tell settlement in relation to other archaeological discoveries in the proximity (B): 1. Morteni-Măgura, 2. Morteni-La Bold, 3. Morteni - Vatra Satului, 4. Morteni-Putul lui Iordache, 5. Morteni-Pădurea Ciocelor, 6. Morteni-La Tufanu lui Radu lui Andrei, 7. Morteni-Murte, 8. Morteni-La Crevedia.. The maps were created using QGIS 3.20.1-Odense (https://qgis.org) using ESRI as basemap (A) and the Cartographic Server for Cultural Heritage managed by the National Heritage Institute of Romania (https://map.cimec.ro/Mapserver/?layer=ran&cod=68262.02) (B).

P. Diaconescu, an archaeologist from the Dâmboviţa County Museum, conducted surveys at the Morteni tell site in 1975 and initiated excavations in 1976 (test pit) and 1978 (archaeological campaign). The 1978 investigations utilised a single trench (24 × 2.5 m) to create a cross-section from the tell’s central high point, sloping southeastward (Supplementary Figure 1A). This main trench was supplemented with two smaller trenches, bringing the total excavated area to 73.5 m^2^. The stratigraphic investigations revealed three primary occupation levels, each with distinct cultural attributions and features, all belonging to the Chalcolithic period (Figure 2). The first two levels (I and II) were attributed to the Gumelniţa culture, phases A2 and B1 (c. 4400–3900 cal BC), while the third level (III) was associated with the Brăteşti culture (c. 3900–3700 cal BC). Additionally, modern pits were identified in the uppermost layers, just beneath the topsoil (Figure 2). According to Diaconescu’s records, the anthropic deposit of the tell settlement extended to a depth of −2.60 m (Diaconescu 1978-1979; Ilie and Dumitru 2010; Ştefan 2010).

**Figure 2.**
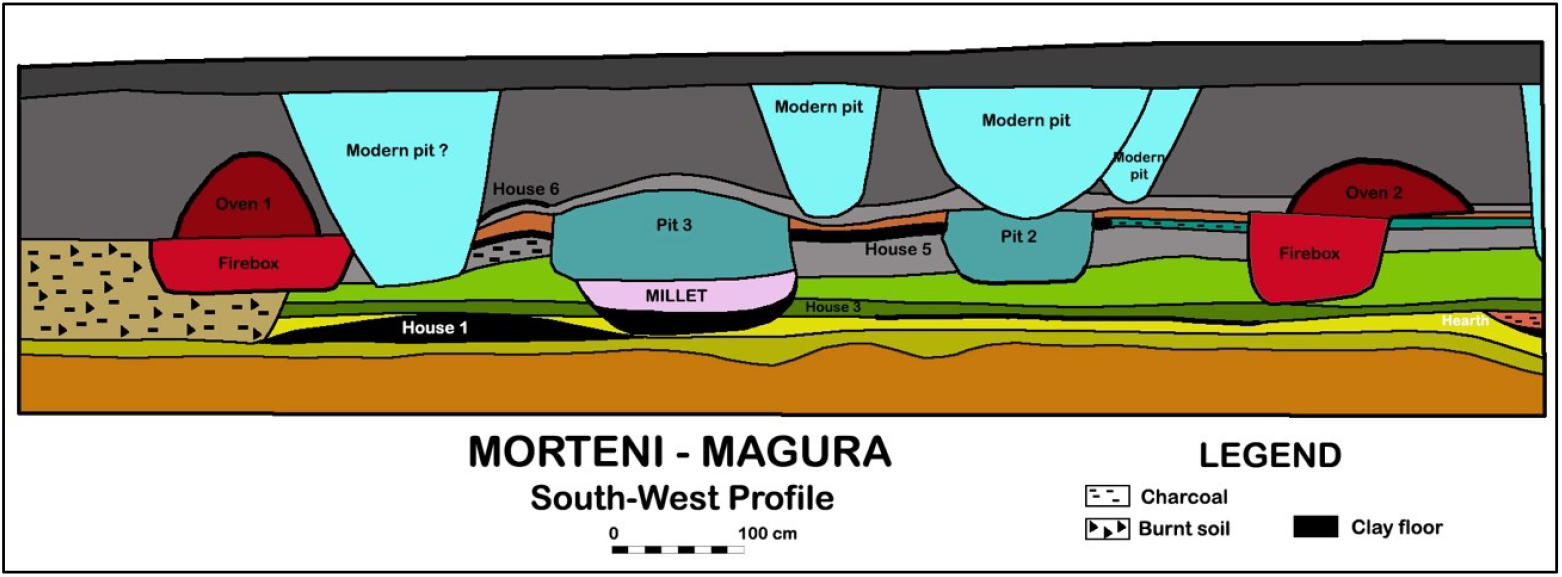
Southwest profile created in 1978 on the Morteni tell, showing the stratigraphic location of Pit 3 and the positioning of the broomcorn millet deposit within it (modified from Diaconescu (1978-1979)).

In 1978, excavations from this site led to a noteworthy discovery that was made in one of the storage pits Pit 3 — also known as the ‘Millet Pit’ (Supplementary Figure 1B) — assigned to the second level (IIb), which belongs to the Gumelniţa culture, phase B1 (c. 4300-3900 cal BC).

Pit 3 had a cylindrical shape with a diameter of 1.80–2.10 m (Supplementary Figure 1B). Like other storage pits, its base was burned and polished to create an impermeable surface, consistent with its function as a storage feature. The lower portion of the pit (Figure 2) contained a substantial amount of botanical material, primarily uncarbonized husks and seeds (approximately 90% of the assemblage). Among these, an extraordinary cache of 150 kg of uncarbonized broomcorn millet, accompanied by other plant species, was identified. This deposit represents one of the largest known prehistoric millet caches in Europe (Diaconescu 1978-1979).

Regrettably, further contextual information about the millet deposit is unavailable, as the research was conducted during a period when data recording and documentation standards were less rigorous. This limitation complicates efforts to fully interpret the circumstances surrounding the storage and deposition of this significant botanical find.

### 2.2. Archaeobotanical remains

The archaeobotanical assemblage from Pit 3 at the Morteni tell site contains 150 kg of remains. It consists of a significant quantity of botanical material, mostly non-carbonised husks and slightly friable grains (around 90%) (Figure 4A, Supplementary Figure 2). The material analysed included a majority of broomcorn millet (88.5%) and smaller proportions of foxtail millet (0.9%), wild buckwheat (*Polygonum convolvulus*, 0.9%), and white goosefoot (*Chenopodium album*, 9.7%) (Cârciumaru 1991; Cârciumaru 1996). The absence of weed seeds and the quantity of stored material suggest deliberate human storage rather than natural deposition by animals or insects. However, there remains some uncertainty about whether the seeds are contemporary with the Gumelniţa culture or originate from a later period (Cârciumaru 1991; Cârciumaru 1996).

Morteni broomcorn husks and grains are not charred, but desiccated, which can be a problem for the preservation of the original isotopic values (DeNiro and Hastorf 1985; Szpak and Chiou 2020), but not for radiocarbon dating.

Although the original assemblage was substantial (approximately 150 kg), only a limited number of husks and grains (n = 20) were available for analysis. This scarcity resulted primarily from the loss of the majority of the assemblage during a reorganization of storage facilities at the Dâmboviţa County Museum in the communist period. Only a small subset was salvaged, from which a significant portion was provided for our study. The 20 specimens obtained were subjected to radiocarbon dating and stable isotope analysis, with morphometric measurements conducted on a subset of these specimens (n = 10). Given that both radiocarbon dating and isotopic analyses involve destructive processes, only the specimens utilized for morphometric measurements remain available. These have been intentionally preserved to facilitate potential future analyses. They should aim to increase the sample size to strengthen the reliability of the results and explore additional analytical methods. Expanding the analytical toolkit is likely to yield further insights, thereby enhancing our understanding of the assemblage’s broader archaeological context and improving interpretative frameworks.

### 2.3. Morphometric analysis

The morphometric analysis was conducted on 10 husks at Babeş-Bolyai University in Cluj-Napoca (Romania), using a binocular magnifier (Olympus Stereo Zoom Binocular Microscope, Olympus Corporation, Tokyo, Japan). The standard morphological examination focused on the shapes observed in dorsal, ventral, lateral, and cross-sectional views, (Figure 4). Measurements of the husks’ length (L), breadth (B), and height (H) were taken using the stereomicroscope. The average, median, and variance were calculated following the methodology of Jacomet (2006).

### 2.4. Radiocarbon dating

Five broomcorn millet remains were selected for radiocarbon dating. The experiments were carried out at the 1 MV Tandetron™ accelerator from RoAMS Laboratory of the “Horia Hulubei” National Institute for Physics and Nuclear Engineering (IFIN-HH), with the analyses conducted at the (Măgurele, Romania). Samples were pretreated following (Sava et al. 2019). Calibration of the resulting data was performed using OxCal v4.4 (Bronk Ramsey 2009) in conjunction with the IntCal20 calibration curve (Reimer et al. 2020).

### 2.5. Stable isotopes analyses

Stable isotope analysis was conducted on five broomcorn millet husks, all of which were intact and may still contain preserved caryopses. To assess potential soil contamination, FTIR-ATR analysis was conducted on MOR-1 using a Bruker Vector 22 at the Unit of Molecular Spectroscopy (UEM) within the Research Support Services (SAI) of the University of A Coruña (UDC). A single measurement was taken, with background subtraction and baseline correction applied via the OPUS software, followed by spectrum normalization. Specific peaks were examined to detect contamination: 870 and 720 cm^−1^ for carbonates; 3300, 1450, and 1085 cm^−1^ for nitrates; and 3690, 1080, and 1010 cm^−1^ for humic acids. As no contamination peaks were identified (Figure 3), the grains were directly analysed using IRMS.

**Figure 3.**
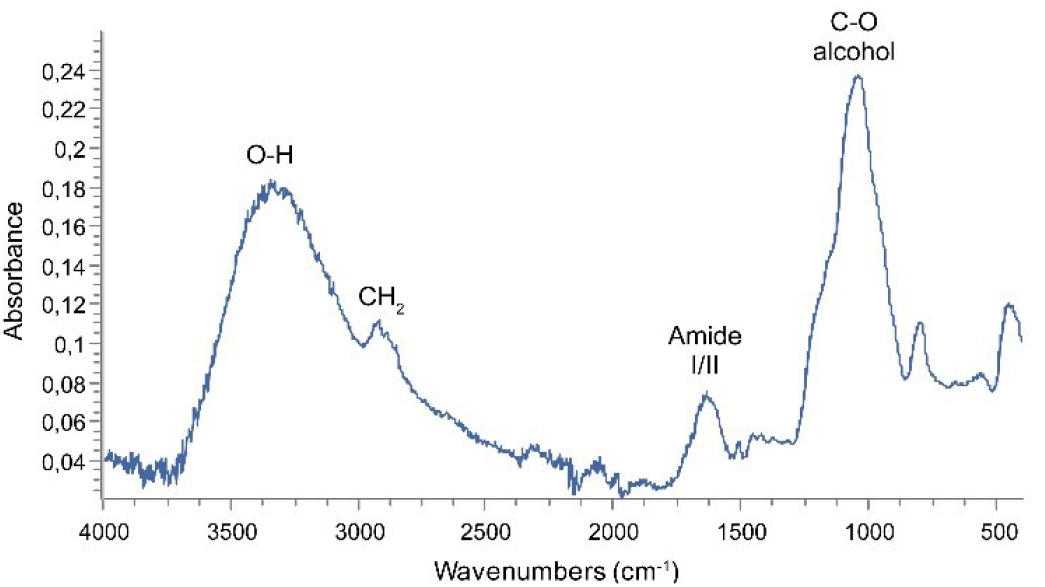
FTIR-ATR spectrum (400–4000 cm^−1^) of the Morteni seed tested (MOR-1). Key peaks are identified following Metcalfe & Mead (2019). Plot produced using the software Spectragryph 1.1.15 (Menges, 2020).

**Figure 4.**
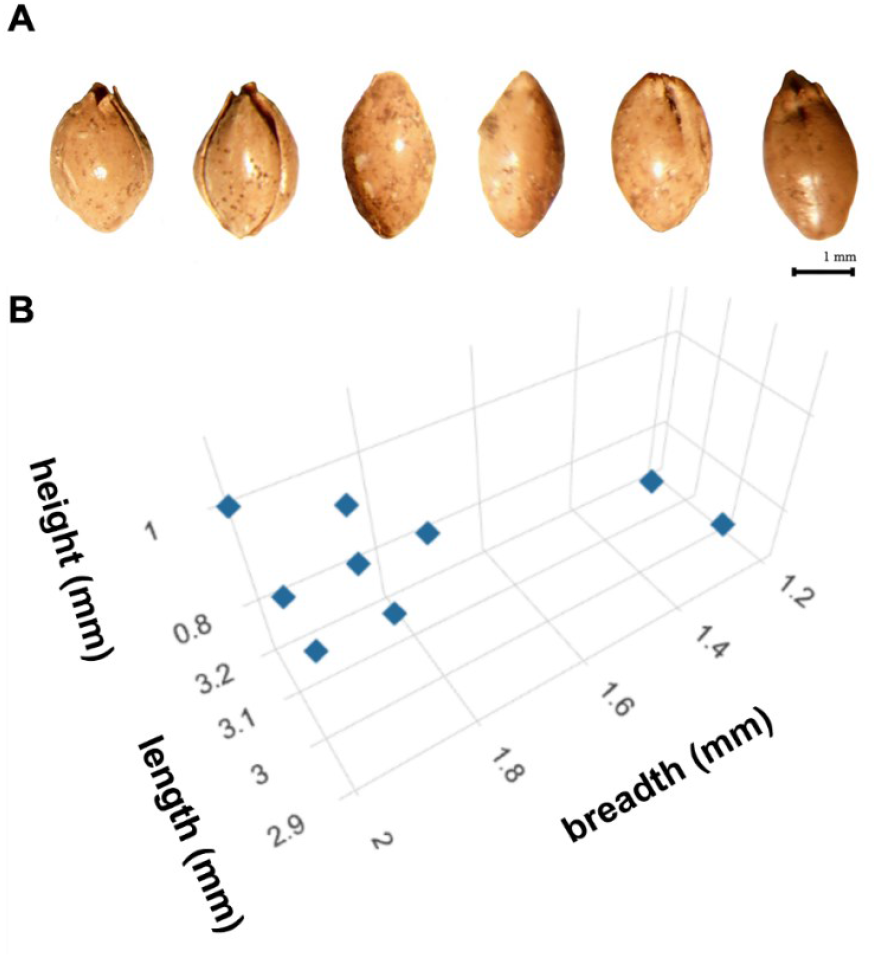
A) Broomcorn millet husks from Morteni tell site. B) 3D plotting of the length, breadth, and height of the ten analysed grains.

The IRMS analysis was performed at the Unit of Instrumental Techniques of Analysis (UTIA) of the SAI (UDC). Samples were combusted and analysed using a DeltaV Advantage (ThermoScientific) coupled with a ConfloIV interface and a Flash IRMS EA IsoLink CNS (ThermoScientific). The method achieved an analytical reproducibility of better than ± 0.2‰ for both carbon and nitrogen. Isotopic compositions were calibrated against international standards (VPDB for carbon and AIR for nitrogen) using secondary standards such as USGS 40, USGS 41a, IAEA-N-1, and others. Internal precision was verified using acetanilide, yielding a standard deviation of ± 0.15‰ across 10 replicates. Each grain was analysed once, with results expressed in delta (δ) notation, which represents the isotopic ratio of the sample relative to the international standard using the equation δX (‰) = [(R_sample_/R_standard_) − 1] × 1000, where X refers to the heavier isotope and R is the ^15^N/^14^N or ^13^C/^12^C ratio.

To evaluate potential diagenetic alterations, given that the samples were desiccated rather than charred, correlations were analysed as recommended by Vaiglova et al. (2023), specifically δ^13^C *vs*. C:N, δ^13^C *vs*. %C, δ^15^N *vs*. C:N, and δ^15^N *vs*. %N.

Statistical analyses and plots were performed using Past 4.14 (Hammer et al. 2001) and RStudio 2023.06.1 (R Core Team 2023), with final edits made in Adobe Illustrator.

## 3. Results

### 3.1. Measurements

The obtained values for length, breadth, and height (Table 1, Supplementary Table 2,) align with the ranges previously reported in the literature for uncharred archaeological husks of broomcorn millet (Liu et al. 2022). These measurements are graphically represented in Figure 4B.

**Table 1.**
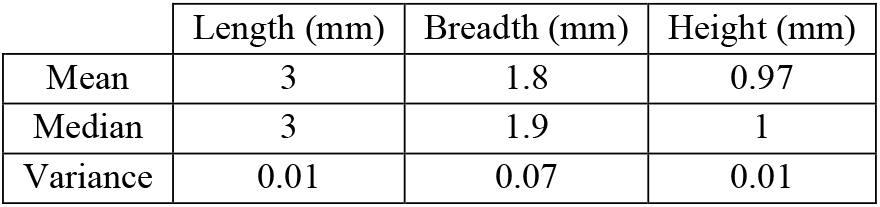
Average, median, and variance of measurements on the analysed millet grains (n = 10)

### 3.2. Radiocarbon dating

The broomcorn millet remains analysed in this study were identified in Chalcolithic context, although radiocarbon dating produced younger results. The sample from Morteni (P222), was dated to 2105 ± 69 BP (128 ± 108 cal BC) (Table 2 and Figure 5), corresponding to the Iron Age rather than the expected Chalcolithic period.

**Table 2.** Radiocarbon results. Calibrated data is expressed by the median + σ, the σ and the 2σ intervals.

| Lab Code | Sample ID | Context | Years BP | Years cal BC (median $\pm \sigma$) | Date calibrated ($\sigma$) | Date calibrated ($2\sigma$) |
| --- | --- | --- | --- | --- | --- | --- |
| RoAMS-5525.2 | P222 | Pit 3 | $2105 \pm 69$ | $128 \pm 108$ | 342 – 1 cal BC (68.3%) | 361 cal BC – 61 cal AD (95.4%) |

**Figure 5.**
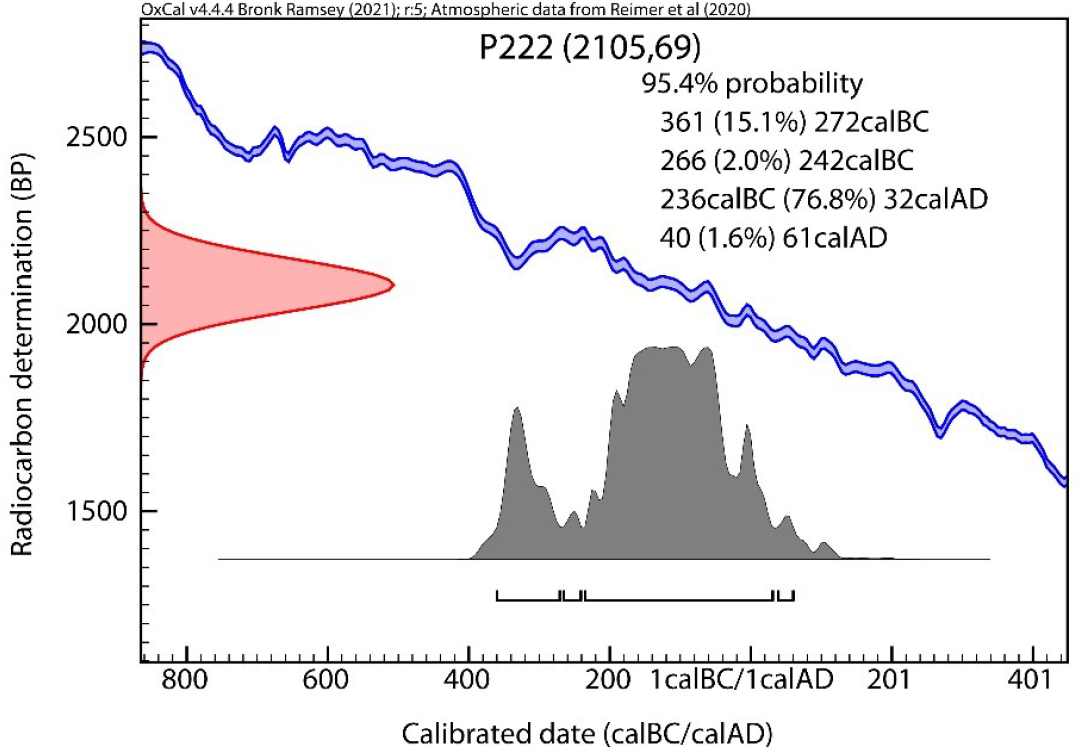
Calibrated date from the broomcorn grains from Morteni.

### 3.3. Stable isotopes analyses

The isotopic results can be found in Table 3. The five husks from Morteni have a mean δ^15^N value of 10.6‰ (range: 9.2 to 11.4‰, SD = 0.85), and a mean δ^13^C value of −12.1‰ (range: −12.6 to −11.8‰, SD = 0.31). The δ^13^C values fall within the typical range for C4 plants (−10 to −16‰) (Bender 1971; O’Leary 1988; Farquhar et al. 1989; Cerling and Harris 1999).

**Table 3.**
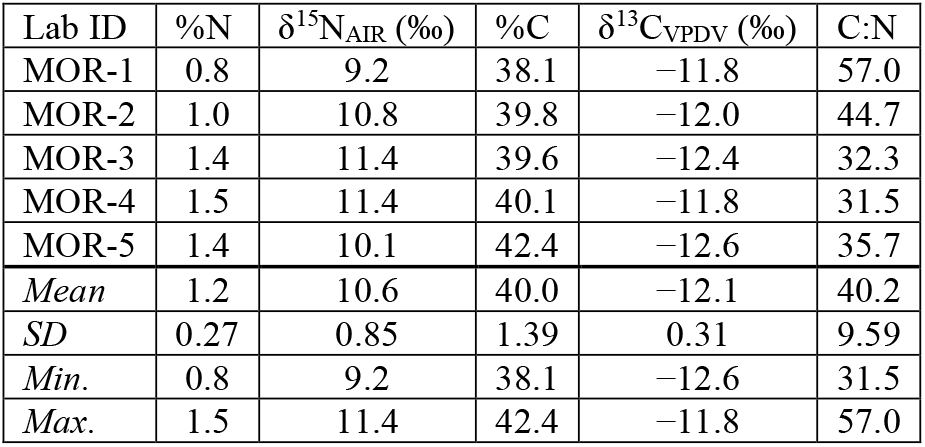
Isotopic results of broomcorn millet from Morteni, and mean, standard deviation, minimum and maximum values of the samples.

As previously mentioned, the archaeobotanical remains are not charred. To evaluate the preservation of the isotopic signals, correlations of δ^13^C *vs*. C:N, δ^13^C *vs*. %C, δ^15^N *vs*. C:N and δ^15^N *vs*. %N were analysed (Figure 6), following the approach outlined by Vaiglova et al., (2023). The relationships between δ^15^N and %N (Figure 6A), δ^15^N and C:N (Figure 6B) and δ^13^C and %C (Figure 6C) indicate that over 50% of the observed variability is explained by the regression models. This suggests that δ^13^C and δ^15^N values are not independent on the amount of carbon and nitrogen of the husks. However, the correlation between δ^13^C and C:N (Figure 6D) is weaker. While these trends are informative, the small sample size limits the statistical robustness of the analysis. It is possible that the inclusion of additional data points could modify the observed relationships and provide a more reliable assessment of isotopic preservation and variability.

**Figure 6.**
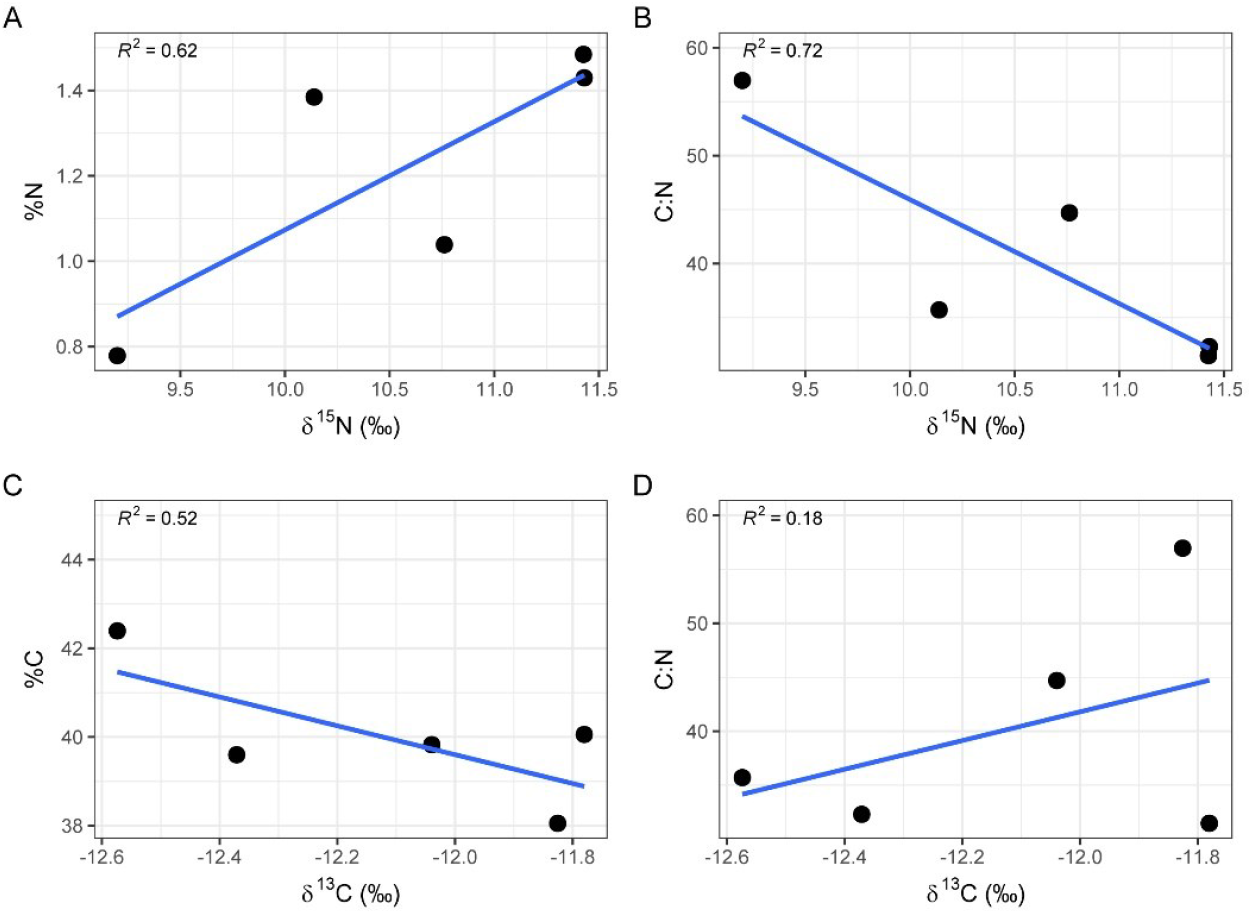
Correlations between the different isotopic parameters of Morteni brooncorn millet grains: A) δ^15^N vs. %N. B) δ^15^N vs. C:N and. C) δ^13^C vs. %C. D) δ^13^C vs. C:N.

## 4. Discussion

The radiocarbon date from Morteni (128 ± 108 cal BC), correspond to the Iron Age, a period when millet consumption had become more widespread (Cârciumaru 1996), and millet-dominated agriculture was prevalent in Eastern Europe (Salova et al. 2024). This finding highlights the critical role of radiocarbon dating, as the pit was initially attributed to the Chalcolithic period, but this radiocarbon date revealed to be a later deposition. A comparable case involved rye grains from a Chalcolithic context from Cuneşti tell site (Romania), which were determined to have been deposited during the Middle Ages (Golea et al. 2023). Additionally, millet grains from Neolithic contexts in Europe have also been shown to be more recent than initially assumed (Motuzaite-Matuzeviciute et al. 2013; Filipović et al. 2020), further emphasising the need for precise dating, especially when dealing with plant remains.

One of the most challenging aspects of this study is explaining the initial chrono-cultural attribution made by the original excavator at Morteni. Archaeological research methods during Romania’s communist period were often constrained by limited funding, restricted access to data, and logistical challenges. The investigation of the Morteni tell began with a field survey in 1975, followed by a small verification trench in 1976, and a single excavation campaign in 1978 (Diaconescu 1978-1979). The southwestern profile reveals the presence of several modern pits in the upper stratigraphy (Figure 2). According to this publication, the uppermost level of the tell (Level III) contained sparse traces of habitation and was heavily disturbed by agricultural activities and modern pits, one of which, though unspecified, was attributed to this final occupation level. Given these disruptions, particularly agricultural activity, it is reasonable to assume that Pit 3, radiocarbon-dated to the Iron Age, originated within Level III (Figure 2) but was heavily impacted, making detailed documentation during the 1978 campaign impossible. Unfortunately, Diaconescu did not provide a detailed inventory of Pit 3, where artefacts are presented in aggregate across the three stratigraphic levels rather than being associated with individual features. Consequently, it remains unclear whether the broomcorn millet grains were the sole contents of this storage pit.

The absence of Iron Age ceramics at the Morteni tell raises questions about the presence of a permanent settlement from this period. Instead, it seems more likely that the elevated position of the Chalcolithic tell in the Neajlov River floodplain was temporarily utilised by Iron Age communities for storage or occasional activities. The tell, rising 3–5 m above the current ground level near a tributary, offered a strategic location for such uses. Similar practices have been documented at contemporaneous sites, such as the Cuneşti tell (Golea et al. 2023), where prehistoric mounds with altimetric advantages were re-used by later communities in flood-prone areas.

Given this evidence, the Morteni tell appears to have functioned primarily as a storage site for a substantial quantity of broomcorn millet during the Iron Age, most likely in its unprocessed form, as suggested by the presence of intact husks indicating the grains were not prepared for consumption This raises the question: where were these communities living? An analysis of the Morteni microzone (Figure 1B, Supplementary Figure 1C) identifies several archaeological sites contemporaneous with or later than the Morteni tell (Diaconescu 1978-1979; Diaconescu et al. 1980-1981; Olteanu 2002). The nearby Latène settlement at Morteni-La Crevedia is particularly noteworthy. Located on a high terrace 200 m east of the Crevedia stream, this site, dated to the 2^nd^ - 1^st^ century BC, covered an area of approximately 3 hectares (Diaconescu et al. 1980-1981; Olteanu 2002) and lies only 3.8 km from the Morteni-Măgura tell. Additionally, Geto-Dacian tetradrachms discovered at Morteni-Vatra Satului, just 0.54 km away from the tell (Olteanu 2002), align chronologically with the radiocarbon dates obtained from the broomcorn millet samples (Table 2, Figure 5). Together, these findings support the interpretation of Pit 3 as an Iron Age storage feature on the Morteni-Măgura tell.

The fact that the broomcorn millet remains from Morteni are desiccated presents a challenge in preserving their original isotopic values, as previously commented. DeNiro & Hastorf (1985) found that seeds charred prior to deposition preserve isotopic values comparable to those of modern plants. However, they observed that uncharred plants, particularly desiccated ones, exhibited altered δ^15^N values and showed much greater variability compared to charred or modern samples. Other authors have suggested that uncharred plants should be carefully considered for palaeoenvironmental and palaeodietary research (Metcalfe and Mead 2019), as they can, under certain circumstances, produce reliable isotopic measurements (Szpak and Chiou 2020).

Our correlations (Figure 6), if evaluated following the criteria of Vaiglova et al. (2023), would suggest the presence of diagenetic alterations in the desiccated grains from Morteni. However, Metcalfe and Mead (2019) dismissed such correlations as unacceptable quality-control indicators for uncharred plants, arguing that similar patterns have been observed in modern plant materials.

Teira-Brión et al. (2024) analysed modern millet grains, both fresh and charred. For fresh hulled grains of broomcorn millet (Supplementary Material Table 2 in Teira-Brión et al. (2024)), their data showed a C:N ratio of 31.2 ± 0.82 (30.3–31.8), %N of 1.6 ± 0.02, and %C of 41.9 ± 0.47 (41.4–42.3). In the case of charred grains, both %C and %N were higher, while the C:N ratio was lower.

Three of the samples analysed in this study, MOR-3, MOR-4, and MOR-5, display C:N ratios, %C, and %N values comparable to those reported for fresh grains by Teira-Brión et al. (2024). However, MOR-1 and MOR-2 exhibit lower %N values, resulting in higher C:N ratios. Interestingly, our FTIR-ATR analysis of MOR-1 (Figure 3) does not indicate contamination that could artificially alter the C:N ratio. However, when examining the peaks and comparing them to those described by Metcalfe and Mead (2019), we observe smaller amide peaks (representing the proteinaceous components) relative to carbohydrate peaks. This may reflect either the diagenetic loss of nitrogen-containing compounds, or the analysis of a tissue component inherently richer in carbohydrates and thus poorer in proteins.. This nitrogen loss increases the C:N ratio and reduces %N in this sample. For this reason, the observed correlations should be interpreted with caution, as not all the grains retain the same level of preservation of their original isotopic values. Furthermore, the C:N ratios may indicate that MOR-3, MOR-4, and MOR-5 correspond to complete hulled grains (caryopsis and husk), while MOR-1 and MOR-2 likely represent husks only, as grains typically contain more protein than husks and therefore tend to exhibit higher nitrogen content (Kapustin et al. 2023). Nevertheless, the δ-values across the entire sample are consistent, and the standard deviations for both δ^15^N and δ^13^C are not particularly large, suggesting a degree of reliability despite potential diagenetic alterations.

The δ^15^N values of MOR-3, MOR-4, and MOR-5, with a mean of 11.0‰, indicate that these plants were grown in a heavily manured field, according to modern experiments (Christensen et al. 2022; Yang et al. 2024). This, combined with the large size of the assemblage, highlights the significant role of broomcorn millet in Iron Age agriculture in Romania. Interestingly, our morphometric analysis reveals two distinct groups regarding the size, a pattern also observed in Liu et al. (2022). These authors associate this differentiation with variations in δ^15^N, noting that larger grains exhibit higher values, with a difference of just over 1‰. Although we did not measure the size of the grains subjected to isotopic analysis, we did observe a δ^15^N difference of 1.3‰ between MOR-5 and MOR-3/4. These differences in size and isotopic values could simply be attributed to variations in manuring, as the similarity in isotopic values suggests that these grains were likely cultivated in the same field during the same year.

The archaeobotanical assemblage of Pit 3 from Morteni, weighing 150 kg, stands as one of the largest broomcorn millet assemblages discovered in a comparable context. This raises critical questions about the number of individuals such a deposit could sustain and the intended purpose behind this substantial accumulation. Although broomcorn millet is not the only species found, it represents 88.5% of the total, which would equate to 132.75 kg and a total of approximately 147 million grains (Supplementary Texts 1 and 2). This quantity of grain could have sustained 3 nuclear families of 4–5 members or a single large, multigenerational household unit for a year (Supplementary Text 2). Combined with the similarity in isotopic data, this could suggest that the broomcorn millet deposit originated from a single harvest, grown in the same field during the same year.

## 5. Conclusions

The radiocarbon dating of the Morteni millet deposit, which dates it to the Iron Age (128 ± 108), highlights the critical importance of precise dating methods, correcting its earlier misattribution to the Chalcolithic period. This finding aligns with patterns observed at other sites (e.g., Cuneşti, Măgura-Buduiasca), where plant remains initially attributed to prehistoric contexts were subsequently found to be significantly more recent. Furthermore, the Morteni tell appears to have functioned as a temporary storage site rather than a settlement during the Iron Age, with its elevated position providing a strategic advantage within the floodplain landscape.

The comparison of our %C, %N, and C:N values with those of fresh broomcorn millet, combined with FTIR-ATR analysis, reveals an uneven preservation of the grains. Specifically, MOR-1 and MOR-2 exhibit a loss of nitrogen-containing compounds, as evidenced by reduced amide peaks in the FTIR analysis of MOR-1, while MOR-3, MOR-4, and MOR-5 show values similar to those of fresh grains. Therefore, the δ-values of MOR-3, MOR-4, and MOR-5 can be considered reliable, whereas those of MOR-1 and MOR-2 probably are not.

Broomcorn millet during this period is one of the most important crops in Eastern Europe, and also in Romania, as confirmed by this study. The metric analysis reveals two potential groups, likely the result of an uneven application of the already abundant manuring of the fields. The data seem to suggest that the deposit originated from a single harvest and could correspond to the granary of one family.

Despite the limited size of the sample set, the data generated provide meaningful new insights. Expanding the sample size and analytical methods remains essential to enhance the robustness of findings and to refine the archaeological interpretation.

Our study highlights the critical importance of radiocarbon dating for archaeobotanical remains, especially those originating from older excavations and published with an incorrect chrono-cultural attributions. Correcting archaeological data already introduced into scientific circulation is both a necessity and an obligation in order to understand better the past realities, a task we have also sought to address in the pages of this study.

## 6. Acknowledgments

The authors want to thank to Aurora Grandal d’Anglade for the access to the laboratory of Molecular Palaeontology of the University Institute of Geology (IUX) of the University of A Coruña (Spain). We also thank Theodor Ignat (Bucharest Municipality Museum, Romania) for his support in processing some of the images and Mihai Florea (National Museum of Romanian History, Romania) for providing additional cartographic data related to the Morteni site. A. García-Vázquez was supported by a Research Fellowship for Visiting Professors within the Research Institute of the University of Bucharest (ICUB) under the project “Stable isotopes of archaeological samples from Romania” (810/16.01.2023) and the program (PNRR-III-C9-2023-I8) of Romanian MCID (64/30.07.2023). Mihaela Golea wish to thank the ‘Vasile Pârvan’ Archaeological Institute and the School of Advance Studies of the Romanian Academy for supporting her PhD. The radiocarbon dating was supported under Research Programme Partnership in Priority Areas PNII MEN-UEFISCDI, contract PN 23210102 and PN 23210201. Radiocarbon dates were supported by the Romanian Government Programme through the National Programme for Infrastructure of National Interest (IOSIN). The Open Access Fee for publication was supported by the University of Bucharest (Romania). Last but not least, the team from the University of Bucharest would like to thank the Vice-Rector for Research of our university, Professor Carmen Chifiriuc, for her support in our research endeavour.

