## Supplementary Materials for "Old Collections, New Perspectives: Analytical Reassessment of One of Europe’s Biggest Archaeological Broomcorn Millet Deposits (Romania)"

#### Supplementary Figure 1

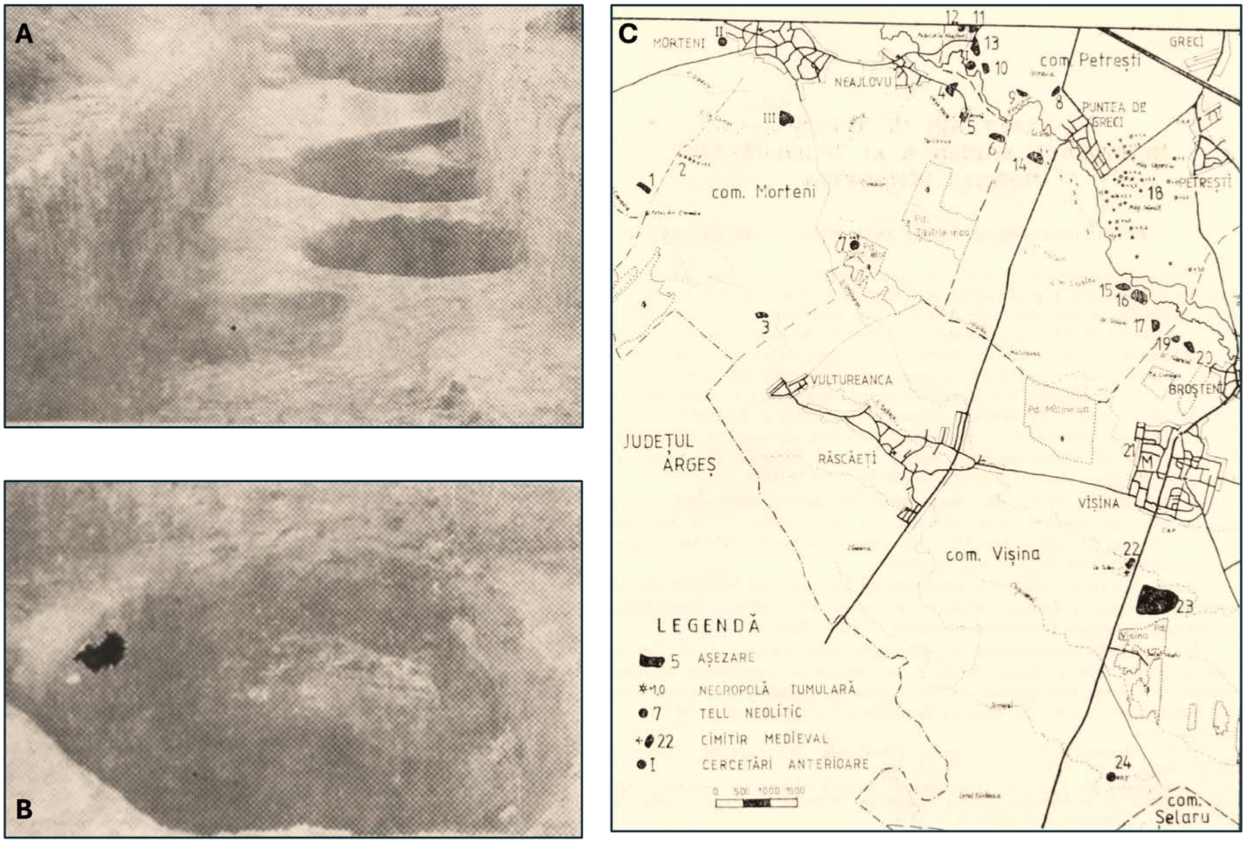

*Supplementary Figure 1. Morteni tell site: A. Image of the 1978 section; B. Image of Pit 3, also known as the "Millet Pit" (after Diaconescu, 1978-1979); C. Map of the surface surveys conducted in the area of the Morteni tell in 1981, showing the location of other archaeological sites (after Diaconescu et al. 1980-1981).*

### Supplementary Figure 2

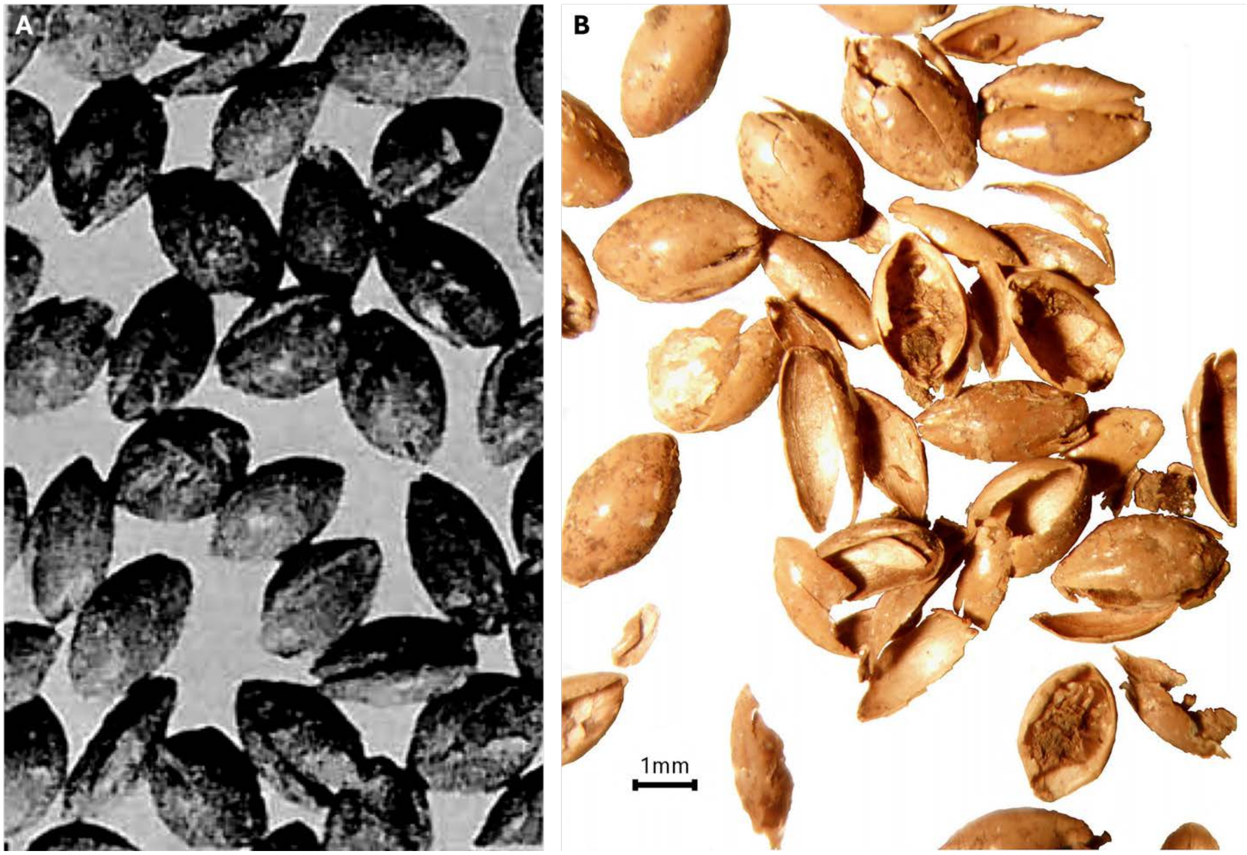

*Supplementary Figure 2. Broomcorn millet assemblage from Morteni: (A) grains photographed during the initial carpological analysis (after Cârciumaru, 1996) and (B) grains during the re-evaluation in 2023.*

### Supplementary Table 1

*Supplementary Table 1. Examples of millet finds in Romania, together with the quantity, chronological period of the site and calibrated radiocarbon data . The radiocarbon data were calibrated using OxCal. v4.4* (Bronk Ramsey 2009) *and the IntCal20 atmospheric curve* (Reimer et al. 2020).

| Site | Chronocultural association | Quantity | Radiocarbon dates | Reference |
|---|---|---|---|---|
| Cârcea | Neolithic | 1 | - | Cârciumaru (1996) |
| Limba | Neolithic | 1 | - | Daisă-Ciută et al. (2004) |
| Măgura Buduiasca | Neolithic | 4 | 1436 - 1263 cal BC<br>1440 - 1623 cal AD | Motuzaite-Matuzeviciute et al. (2013) |
| Miercurea-Sibiului | Neolithic | few | 364 - 121 cal BC | Filipović et al. (2020) |
| Bordușani | Chalcolithic | 7 | - | Monah (2007) |
| Baia-În Muchie | Chalcolithic - Iron Age | 97 | 166–335 cal AD<br>241–352 cal AD<br>247–404 cal AD<br>250–413 cal AD | An et al. (2025) |
| Cheile Turzii | Chalcolithic | 1 | - | Ciută (2006) |
| Dobrovăț | Chalcolithic | 117 | 1442 - 1285 cal BC<br>1114 - 924 cal BC<br>1002 - 821 cal BC<br>1007 - 891 cal BC<br>1018 - 895 cal BC<br>932 - 813 cal BC<br>902 - 803 cal BC<br>805 - 747 cal BC<br>124 - 250 cal AD<br>890 - 1020 cal AD | An et al. (2025) |
| Gumelnița | Chalcolithic | 6 | - | Lazăr et al. (2020) |
| Poduri- Dealul Ghindaru | Chalcolithic | | - | Monah and Monah (2005) |
| Văleni | Chalcolithic | 4 | - | Cârciumaru (1996) |
| Odaia Turcului | Broze Age | Thousands | - | Cârciumaru (1996) |
| Cârlomănești | Bronze Age | Several thousands | - | Cârciumaru (1996) |
| Cornești | Bronze Age | | 1396 - 1227 cal BC<br>1398 - 1226 cal BC<br>1407 - 1222 cal BC<br>1410 - 1265 cal BC<br>1417 - 1284 cal BC<br>1418 - 1284 cal BC | Filipović et al. (2020) |
| Sânzieni | Bronze Age | 2600 | - | Cârciumaru (1996) |
| Teleac | Bronze Age | 1 | 983 - 822 cal BC | Filipović et al. (2020) |
| Cândești | Bronze Age / Iron Age | 7 | - | Cârciumaru (1983) |
| Babadag | Iron Age | 4 | - | Cârciumaru (1996) |
| Tășad | Iron Age | 2 | - | Cârciumaru (1983) |
| Brad | Iron Age | 1294 | - | Cârciumaru (1983) |
| Bâzdâna | Iron Age | 27.5% | - | Cârciumaru (1996) |
| Satu-Nou | Iron Age | > 5 | - | Cârciumaru (1996) |
| Socu | Iron Age | Mass finds | - | Cârciumaru (1996) |
| Grădiștea | Iron Age / Antiquity | 28 | - | Cârciumaru (1996) |
| Bărboși | Antiquity | 100 g | - | Cârciumaru (1996) |
| Biharia | Antiquity | 700 | - | Cârciumaru (1996) |
| Căpâlna | Antiquity | 5 | - | Cârciumaru (1996) |
| Grădiștea Muncelului | Antiquity | >57 | - | Cârciumaru (1996) |
| Hinova | Antiquity | > 500 | - | Cârciumaru (1996) |
| Ocnița | Antiquity | 300 g | - | Cârciumaru (1996) |

| | | | | |
|---|---|---|---|---|
| Piscul Crăsani | Antiquity | >1 | - | Cârciumaru (1996) |
| Popești | Antiquity | > 800 | - | Cârciumaru (1996) |
| Spineni | Antiquity | 4 | - | Cârciumaru (1996) |
| Dinogeția | Late Antiquity | 1 | - | Cârciumaru and Dincă (2000) |
| Topraichioi | Late Antiquity | 300 g | - | Cârciumaru (1996) |
| Dinogeția | Late Antiquity / Middle Ages | Several thousands | - | Cârciumaru and Dincă (2000) |
| Fundu Herții | Middle Ages | Mass finds | - | Cârciumaru and Dincă (2000) |
| Izvoare | Middle Ages | Mass finds | - | Cârciumaru and Dincă (2000) |
| Nufărul | Middle Ages | 21 | - | Cârciumaru and Dincă (2000) |
| Păcuiul lui Soare | Middle Ages | Several thousands | - | Cârciumaru and Dincă (2000) |
| Uivar | Middle Ages | 10 | - | Fischer and Rösch (2004) |
| Dîngeni | Late Middle Ages | Several thousands | - | Cârciumaru and Dincă (2000) |

#### Supplementary Table 2

*Supplementary Table 2. Length, breadth, and height of the ten analysed millet husks from Morteni tell site.*

| Millet caryopse | Length (L) | Breadth (B) | Height (H) |
|---|---|---|---|
| 1 | 3 | 1.9 | 1.1 |
| 2 | 3 | 2 | 1 |
| 3 | 3 | 1.9 | 1 |
| 4 | 3 | 1.9 | 1 |
| 5 | 3 | 1.8 | 1 |
| 6 | 2.9 | 2 | 1 |
| 7 | 2.9 | 1.9 | 1 |
| 8 | 3.2 | 2 | 1 |
| 9 | 3 | 1.2 | 0.7 |
| 10 | 3 | 1.4 | 0.9 |
| *Mean* | 3 | 1.8 | 0.97 |
| *Median* | 3 | 1.9 | 1 |
| *Variance* | 0.01 | 0.07 | 0.01 |

#### Supplementary Text 1

To estimate the total number of broomcorn millet grains in Pit 3, it was determined that they constitute 88.5% of the assemblage. The average weight of individual husks, calculated as 0.902 mg (n = 4), is detailed in Supplementary Table 3. With the total weight of the assemblage being 150 kg, the broomcorn millet portion accounts for 132.75 kg. Using this value, the estimated number of husks is approximately 147 million (147,132,170 grains), as determined through Equation 1.

*Supplementary Table 3. Weight in mg of the broomcorn millet grains analysed isotopically. MOR-1 was excluded from the mean calculation as a portion was used for FTIR-ATR analysis.*

| **Sample ID** | **Weight (mg)** |
|---|---|
| MOR-1 | 0.515 |
| MOR-2 | 0.807 |
| MOR-3 | 0.902 |
| MOR-4 | 1.110 |
| MOR-5 | 0.790 |
| *Mean* | 0.902 |

Equation 1: $Nº\ of\ broomcorn\ husks = \frac{132750000\ mg}{0.902\ mg} = 147132169{,}6\ husks$

#### Supplementary Text 2

To estimate the number of individuals that could have been sustained by the grain recovered from Pit 3, it is first necessary to account for the inedible hull, as broomcorn millet husks are not consumed by humans. Using the weights of husked and dehusked uncharred grains reported in Supplementary Material Table 1 of Teira-Brión et al. (2024) (n = 300), it was determined that the hull of *Panicum miliaceum* constitutes, on average, 16.43% of the total grain weight.

The precise ratio of preserved husks to intact grains remains unclear; thus, we adopt a conservative approach, assuming that the preserved material consists entirely of husks. Under this assumption, the average measured weight of the husks (0.902 mg) would represent 16.43% of the original grain weight, suggesting that a complete, unprocessed grain would have weighed approximately 5.49 mg—comparable to the values reported by Teira-Brión et al. (2024) for uncharred, husked grains (6.27 mg). Accordingly, the 132.75 kg of broomcorn millet recovered from Pit 3 would originally correspond to approximately 807.97 kg of hulled grain, yielding around 675.22 kg of edible seed.

Based on the recommended daily caloric intake of 2,000 kcal for adults, older children, pregnant and lactating women, and considering that 100 g of broomcorn millet provides 378 kcal (Das et al. 2019), meeting this energy requirement would require a daily consumption of 529 g of millet. Under these conditions, the 675.22 kg of grain could provide approximately 1,276 full daily rations—enough to sustain 3–4 individuals for one year. However, it is unlikely that millet constituted the sole dietary component for these communities. The inclusion of other foodstuffs such as legumes, vegetables, and animal products would have reduced the reliance on millet, thereby extending the number of rations derived from the stored grain. For instance, assuming a hypothetical daily millet intake of 50 g per person, the deposit could yield around 13,504 rations—sufficient to sustain approximately 37 individuals for one year. Alternatively, at a daily intake of 125 g per person, the grain could have fed 14–15 individuals annually, which is consistent with the idea that this deposit may have been intended to supply either three nuclear families of 4–5 members or a single large, multigenerational household unit. However, it must be noted that these estimations are likely to represent maximum values, as the actual ratio of preserved husks to intact grains remains unknown. The calculations are based on the assumption that the entire preserved weight corresponds to husks, and our measured grain weights derive exclusively from husk fragments, not from hulled grains.
